# AgeML: Age modeling with Machine Learning

**DOI:** 10.1101/2024.05.02.592130

**Authors:** Jorge Garcia Condado, Iñigo Tellaetxe Elorriaga, Jesus M. Cortes, Asier Erramuzpe

## Abstract

An approach to age modeling involves the supervised prediction of age using machine learning from subject features. The derived age metrics are used to study the relationship between healthy and pathological aging in multiple body systems, as well as the interactions between them. We lack a standard for this type of age modeling. In this work we developed AgeML, an OpenSource software for age-prediction from any type of tabular clinical data following well-established and tested methodologies. The objective is to set standards for reproducibility and standardization of reporting in supervised age modeling tasks. AgeML does age modeling, calculates age deltas, the difference between predicted and chronological age, measures correlations between age deltas and factors, visualizes differences in age deltas of different clinical populations and classifies clinical populations based on age deltas. With this software we are able to reproduce published work and unveil novel relationships between body organs and polygenetic risk scores. AgeML is age modeling made easy for standardization and reproducibility.

## I. Introduction

Age modeling can be implemented using supervised machine learning regression in which clinical, behavioral, cognitive or brain/body morphological features of a subject are used to predict their chronological age [1]. Age models can be built using body organ specific features to create organ specific models, or organ ages [1]. Models have been built for many organs such as the brain [2]–[4], heart [5], eye [6], but also from cognitive performance [7]. These models can integrate information from a single source of data or from multiple sources, thereby facilitating the development of multi-modal age models. The emergence of such age models is attributed to the accessibility of large-scale healthcare databases in the recent decade, exemplified by the UKBiobank [8].

Researchers are focused on elucidating the age gap or delta, the discrepancy between the predicted age and the chronological age of a subject. Age models are trained on two different types of populations: healthy participants devoid of chronic disorders, or on a sample representative of the general population. These deltas, representing the residuals in the regression task, have been shown to correlate with lifestyle factors. Participants engaging in unhealthy lifestyle practices, including smoking, excessive alcohol consumption, or sedentary behavior, exhibit elevated deltas [1]. Age deltas have also been recently associated to specific genetic mutations [9]. The variation in age deltas across organs at different time points, and their multi-variate integration, can help us understand large-scale physiological multi-organ networks [1], of special relevance for the field of network physiology [10]. Participants diagnosed with diseases exhibit higher deltas in specific organs; for example, variations in brain age deltas can distinguish individuals with a neurodegenerative disease such as Alzheimer’s disease from healthy controls [11].

These deltas have two key applications. First, they can be used for screening and identifying outliers that may require further clinical follow-up. For instance, previous studies have demonstrated that brain age deltas are correlated with the time to conversion for Alzheimer’s disease, providing a quantifiable measure of risk [7]. Second, deltas can help quantify deviations from healthy aging and be included as covariates in research, allowing for the separation of aging-related effects from pathological processes [12], [13]. However, from the practical perspective of implementing models and calculating deltas, there still exist many problems and questions in this emerging multidisciplinary field, particularly concerning the implementation, standardization, and reproducibility across institutions.

AgeML is an OpenSource Python package that makes age modeling with machine learning easy. AgeML has three main goals: standardization, reproducibility and easy access. Standardization is the first element necessary to enable the comparison of studies. Reproducibility is crucial for fostering trust within the community and ensuring the consistency of results. It is vital to provide an easy start for individuals interested in age modeling, particularly for clinicians with potentially less theoretical backgrounds in machine learning. We believe that AgeML could serve as a foundational platform for addressing the research questions within the field, led by developers and researchers themselves.

## II. Related Work

Previous works on machine learning-based age modeling have focused on the derived biomarker called the age delta, the difference between the chronological and predicted age. Specifically, understanding if these deltas can be used as a proxy for deviations from a healthy aging trajectory. Research has focused on whether these deviations come from a sudden change, an increase in the delta at a specific time point, or an accumulation of small changes leading to a higher delta over time [14]. This age delta can be used for a classification task downstream. A debate exists regarding whether research should prioritize minimizing the error in age estimation tasks or, alternatively, focus on developing models that lead to deltas that are generalizable for subsequent classification tasks [15]– [17], ie., differentiating between individuals at higher risk of developing a pathological condition. There is also large variability between reported average deltas of disease groups across different studies [18]. Caution is essential in selecting training data, as it often comes from white populations. This could lead to possible biases, such as an increase in age prediction error in controls. This has been observed when applying models trained on white populations to non-white populations [19], [20]. Similar biases have also been found when applying a model across different age ranges than those it was trained in [21]. To tackle all these open questions, standardization in age modeling procedures is needed.

A previous effort aimed at developing a tool specific for brain age modeling is the Brain Age Standardized Evaluation (BASE) initiative [22]. BASE focuses on age modeling from brain magnetic resonance images (MRI), encompassing various scenarios such as scanning at multi-site, generalizing to unseen sites, test-retest, and longitudinal data. It is well known that MRI images can vary greatly between sites and even between images of the same subject taken with the same machine [23]–[25]. It also employs an evaluation protocol involving repeated training of brain age prediction models, assessing performance metrics for accuracy and robustness, to ultimately ensure reproducibility.

BASE is limited to working with neuroimaging data and produces BrainAge models, whereas AgeML operates with tabular data, allowing it to generate models for various organs beyond the brain. For example, while AgeML can produce BrainAge models from brain volumetric tabular data, it can also create models for the heart using routine clinical heart evaluations. Importantly, images are often more expensive, time-consuming, and less accessible compared to tabular data from routine clinical practice due to anonymity concerns. Additionally, images typically require extensive preprocessing compared to tabular data. Age models based on imaging also demand more complex architectures and larger datasets, making them harder to train compared to tabular data-based models. However, one limitation of AgeML is the need to convert all data into tabular form, which could potentially result in a loss of information.

AgeML expands upon many of the functionalities of BASE, enabling multi-modal inputs of different organ features, which aligns with a more contemporary approach of system medicine viewing the body as a network of interacting organs [10]. This makes AgeML more accessible to a broader research community. Moreover, AgeML is open-source developed with transparency to ensure that standards are led by the community. Additionally, AgeML is specifically focused on analysis procedures for understanding clinical populations, thereby offering added value to the medical research community. There is a need to understand model variability [17] and for this to be done properly we need to ensure standardization and reproducibility across all different organ age models for different pathologies.

## III. Data

The UKBiobank database [8] was used for this project (project ID 79105). Overall, the UKBiobank consists of a total of 502,504 individuals, of which 229,122 are males. These participants, aged between 37 and 73 years during recruitment (2006–2010), underwent thorough assessments, including questionnaires, physical examinations, blood and urine sample analyses, and genome-wide genotyping. The extensive data collection occurred across 22 assessment centers located throughout the United Kingdom.

### Body phenotypes

We created age models using 78 different body phenotypes, which align with those in the work of Ye Tian et al. 2023 [1]. A breakdown of these body phenotypes can be found in Supplementary Table S1 of Ye Tian et al. 2023 [1]. The different body phenotypes were grouped into seven distinct organ systems: Musculoskeletal, Cardiovascular, Pulmonary, Immune, Renal, Hepatic, and Metabolic. The organs each phenotype belongs to can be found under the Body age phenotypes subsection of Methods in Ye Tian et al. 2023 [1]. These groups were used to create distinct age models for each organ system. The 78 distinct phenotypes were also considered as features to create a unified age model, referred to as Body.

### Clinical categories

We established them using ICD-10, ICD-9 codes and self-reported pathologies, translated into clinical categories by using the Clinical Classifications Software Refined [26]. Data was used based only on first occurrence labels. The distinction between chronic and non-chronic participants was established using the Chronic Condition Indicator Refined [27], resulting in a healthy population of N=6,388. Eight clinical categories were finally selected, concerning the organ systems previously defined: blood diseases (BLD), circulatory system diseases (CIR), endocrine, nutritional and metabolic diseases (END), mental, behavioral and neurodevelopmental disorders (MBD), Neoplasms (NEO), diseases of the musculoskeletal system (MUS), diseases of the nervous system (NVS), and respiratory system diseases (RSP). Participants with multiple comorbidities were excluded to mitigate potential mixed effects.

### Polygenetic Risk Scores (PRS)

To investigate the relationship between organ deltas and genetics, if any, a factor correlation analysis was carried out with PRS derived from the genetic data of the UKBiobank. Two types of PRS were used: Standard PRS and Enhanced PRS [28]. Standard PRS are sourced from external UKBiobank GWAS data, while Enhanced PRS are derived from a combination of external and internal UK Biobank data. Notably, Enhanced PRS encompass a larger number of categories (53) compared to Standard PRS (39). Standard PRS were available for 6,323 out of the 6,388 participants, whereas Enhanced PRS were only accessible for 1,234 of these individuals. Prior to conducting the factor correlation analysis, the PRS were adjusted for the covariates recruitment site, sex, and ethnicity.

Additional information regarding the total number of participants per category and their age distributions can be found in Table I.

**TABLE I.** Age distribution summary of the clinical categories.

| CLINICAL CATEGORY | $\mu_{age} \pm \sigma_{age}$ (YEARS) | AGE RANGE (YEARS) | N (PARTICIPANTS) |
| --- | --- | --- | --- |
| CN | 51.39 ± 7.31 | [40-70] | 6,388 |
| CIR | 56.00 ± 7.90 | [40-70] | 3,907 |
| END | 55.43 ± 7.78 | [40-70] | 1,616 |
| MBD | 50.86 ± 7.04 | [40-70] | 1,184 |
| MUS | 54.11 ± 7.67 | [39-70] | 5,603 |
| NVS | 53.10 ± 7.63 | [39-70] | 983 |
| RSP | 51.41 ± 7.57 | [40-70] | 4,376 |
| NEO | 56.09 ± 7.75 | [40-70] | 2,215 |
| BLD | 51.76 ± 7.33 | [40-69] | 338 |

CN: Healthy Controls, CIR: Circulatory system disease, MBD: neurodevelopmental disorders, MUS: Musculoskeletal system disorders, RSP: respiratory system diseases, NEO: neoplasms, BLD: blood diseases

## IV. Methods

AgeML is developed as a Python package readily available to install through PyPI [29]. An overview of all AgeML functionalities can be seen in Fig. 1. There are three main strategies to work with AgeML. The first one is a simple command line interface that can be launched after installation by using the command `ageml`. This interface interactively asks for the required individual inputs at each different stage. The second strategy is by using the four available commands, each for a different pipeline: `model_age` for age modeling, `factor_correlation` to study the relationships between deltas and clinical outcomes, `clinical_groups` to study the deltas of different clinical groups and `clinical_classify` to use deltas as predictors in a classification task. Lastly, `ageml` can be used as an importable Python library, allowing for the implementation of custom functionality. The Open Source code can be found in GitHub: https://github.com/compneurobilbao/ageml.

**Fig. 1.**
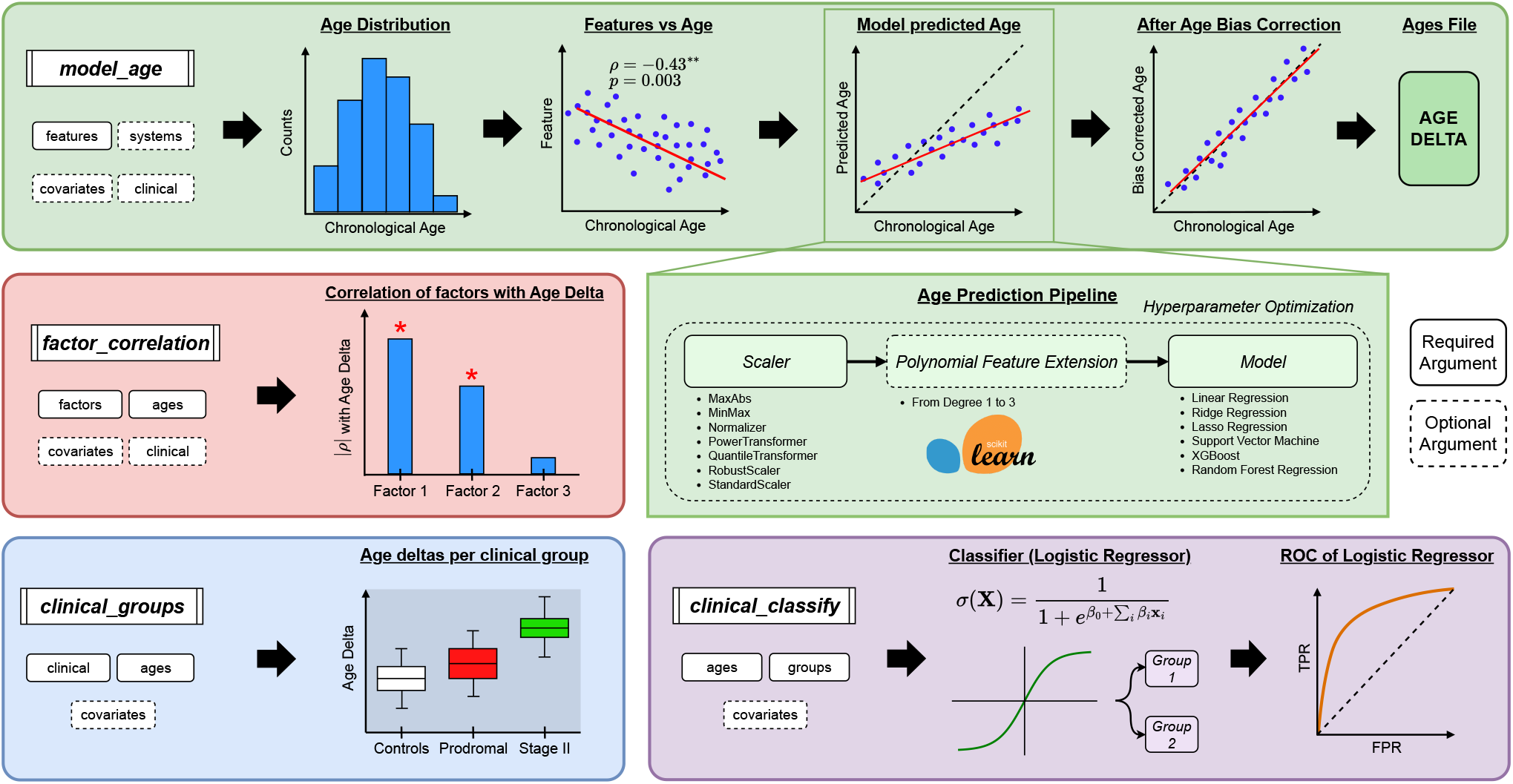
Age modeling with Machine Learning (AgeML), general overview, and major capabilities. *Green panels:* the `model age` module requires only a features file to operate, which should contain the chronological age of the subject and the set of features **x**_***i***_ for each individual. A system file consisting of sets of features (e.g.: a set that encompasses the musculoskeletal system and/or another set referring to renal features) can be provided to train an age prediction model for each specific system. For training a model that distinguishes participants based on a categorical value, a covariates file can be provided, enabling the performance of covariate correction. If a clinical file is provided indicating which participants are healthy controls, the age prediction model is exclusively trained using their features. This pipeline displays the age distribution of the healthy controls, the Pearson correlation (***ρ***) and significance of each feature and chronological age (separated by system and covariate as required), and the predicted age as well as the bias corrected age. Finally, an ages file in CSV format is generated, incorporating the participant’s chronological ages, their predicted uncorrected, corrected ages and the delta, the difference between corrected and chronological age. The age prediction pipeline includes the following components: a scaler selection step, an optional feature extension step, a model selection step, and the availability of optional hyperparameter optimization. *Red panel:* the `factor_correlations` module analyzes the Pearson correlation ***ρ*** strength and significance between the specified numerical factors (e.g.: weekly exercise hours, years of education) and the calculated age delta. Additionally, covariate correction can be applied, and this analysis can be performed for each clinical group as needed. *Blue panel:*, the `clinical_groups` module creates a box plot illustrating the age deltas for each clinical group, reporting the FDR and Bonferroni corrected T-test p-values and T-statistics of all groups. Although not depicted in the figure, the age distributions of all clinical groups are plotted, and a T-test is performed to assess group differences, reporting their p-values. *Purple panel:* The `clinical_classify` module employs Logistic Regression with *n*-fold cross-validation and a defined sigmoid threshold to differentiate between two clinical categories identified in the clinical file. This classification is exclusively based on the age deltas within the ages file. If the ages file comprises multiple systems, individual models are trained for each system, and a comprehensive model encompassing all system-specific age deltas is also trained.

AgeML provides a comprehensive machine learning pipeline for age modeling with four core functions. The first module `model age` (green panel in Fig. 1) predicts age using a features file with optional system-specific modeling. It allows for the training of models on healthy controls and outputting predicted, corrected ages, and the delta (difference between corrected and chronological age) for each of the uploaded groups. The second module `factor correlation` (red panel in Fig. 1) analyzes Pearson correlations between selected numerical factors (e.g., exercise hours) and the age delta, with the option to apply covariate correction. This can be done separately for each clinical group; based only on the control group and then applied to the whole population; or based on the whole population and then applied to it. The third module `clinical groups` (blue panel in Fig. 1) visualizes age deltas across different clinical groups using box plots and performs statistical tests, including FDR and Bonferroni-corrected T-tests, to assess group differences in age deltas. The fourth module `clinical classify` (purple panel in Fig. 1) utilizes Logistic Regression with cross-validation to classify individuals into clinical categories based on age deltas. It supports system-specific and comprehensive models across multiple systems. This structured pipeline allows for flexible age prediction, factor correlation analysis, and clinical group differentiation based on machine learning methods. A toy dataset is provided and concrete examples of use and expected outputs can be found in the Tutorial.

### A. Age modeling

Age modeling can be effectively applied through supervised machine learning regression, where subject features are employed to predict the known chronological age of the subject (*Y*), corresponding to the subject’s birth date. Let *Y* denote the predicted age, and **X** represent the model features. The aim is to find *f* (), our age modeling pipeline, such that:

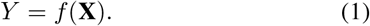

The pipeline consists of three steps: a scaler, a feature extension step, and a model. The available scalers are all imported from the Scikit-learn Python library [30] and include the max absolute scaler, min-max scaler, normalizer, power transformer, quantile transformer, robust scaler, and standard scaler. For feature extension, the polynomial feature extension from Scikit-learn [30] is used. The various models implemented consist of Linear Regression, Ridge, LASSO, Support Vector Regression (SVR), Random Forests (RF) and XGBoost [31].

AgeML contains a features vs age component that offers a visual and statistical check to assess potential correlations between features and chronological age. By computing Pearson’s correlation coefficients, it provides a quick sanity check on whether the features may be informative for age prediction. While Pearson’s correlation only captures linear relationships, visual inspection of the data may reveal non-linear patterns, prompting users to explore models better suited for capturing such relationships. This step ensures a more comprehensive understanding of the feature set and its relevance for accurate age modeling.

The feature extension module is not used as a default. The feature extension is left for modeling non-linear interactions of variables that could help in the downstream age modeling task. Many body variables follow a u-shaped pattern with age increasing until mid age and then decreasing in old age or vice versa. For example, when doing brain age modeling over a wide age range of between 5 to 100 years there are many volumetric features that follow this u-shape [32], [33]. With feature extension volumetric features extended as 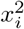 or *x*_*i*_ *× x*_*y*_ become linearly correlated with age. Including the feature extension module in these cases could help to model these relationships in the age modeling task while using linear regression models instead of using more complex models such as SVM to capture this known interaction. Keeping a linear regression model with feature extension can allow for more explainability.

The pipeline is trained by minimizing the mean squared error between the predicted and chronological age. The pipeline undergoes training via cross-validation, with a default of 5-folds. The reported metrics are derived from these cross-validation folds. Users have the option to configure the hyperparameters of the model manually, or they can choose to do hyperparameter tuning. Hyperparameter tuning can be performed by choosing a specific model and optimizing the hyperparameters using cross validation and a grid search with a specified number of points. Alternatively, it can let AgeML choose the best model and preprocessing leveraging the capabilities of the hyperopt-sklearn library [34].

A well-known bias in age modeling is the tendency for younger participants to be assigned higher ages, while older participants are assigned lower ages than their actual ages. This phenomenon is attributed to a regression to the mean problem [21], [35], [36]. This bias can be conceptualized as our best estimate for a subject about whom we have no prior information being the mean age. The predicted age bias can be can be corrected [35], by taking into account the actual participant’s age through fitting the following linear model :

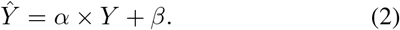

*Ŷ* is the predicted age. The coefficients *α* and *β* correspond to the slope and intercept of the linear model. The training set predictions *Ŷ*_*train*_ and chronological age *Y*_*train*_ is used to fit these coefficients *α* and *β*. Selecting the appropriate method for age bias correction is of paramount importance to prevent the introduction of new biases, such as inflating correlation effects. An alternative method to that used in this study is to use age as a covariate when studying group differences [35]. The fitted *α* and *β* are then used to correct the predictions in a test set to obtain an age bias corrected prediction 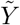 using:

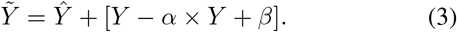

Research in this context typically focused on the age gap or delta (*δ*), which is defined as the difference between the subject’s predicted age after age bias correction and the subject’s chronological age:

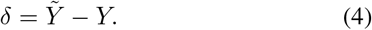

AgeML provides the capability to simultaneously train distinct models for each system defined, thereby obtaining age deltas for each system. This is achieved by specifying which features belong to each system. Additionally, separate models can be trained based on a covariate, such as sex. This would train distinct models for males and females for each system.

Users also have the option to specify which participants are considered as healthy controls for model training purposes. The model is then trained exclusively on these designated participants and subsequently applied to calculate deltas for the remaining participants. The predicted age (*Ŷ*), the predicted age after age bias correction 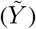 and the age delta (*δ*) are saved and can be used as inputs for further steps or for independent further analysis by the researcher, referred to as the *ages* file. To ensure standardization and reproducibility, after each model is trained, both training and test metrics are stored. These metrics encompass Mean Absolute Error (MAE), Root Mean Square Error (RMSE), *R*^2^, and Pearson’s correlation (*ρ*). Recognizing that MAE and RMSE are strongly influenced by the age range of the training data, an additional metric referred to as the *dummy regressor* (DR) is calculated. The DR is obtained by using the mean age of control subjects as predictor for all subjects and calculating the resulting MAE and RMSE.

### B. Factor Correlation

Previous research has addressed the relationship between the deltas and other external factors such as lifestyle choices or genetics. To visualize and quantify this relations, a Pearson correlation is calculated between each factor (*Z*) and each delta (*δ*) as well as the corresponding p-values. To account for multiple comparisons, p-values can be adjusted using either Bonferroni or False Discovery Rate (FDR) corrections. Additionally, users have the option to provide covariates (*W*) and remove covariate effects from the factors prior to conducting the correlation analysis. The covariate correction is calculated as the coefficients in the maximum likelihood solution of a general linear model, ie.,

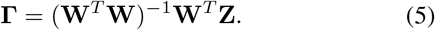

As previously mentioned, **Γ** can be calculated based on only healthy controls, the whole population, or for each clinical group. Providing different covariate correction schemes enables accounting for complex covariate effects across diverse clinical populations, as their characteristics and their relation with covariates can vary significantly. The participant’s covariate-corrected factors are then given by:

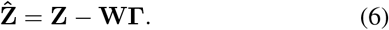

The correlation analysis between factors and deltas is subsequently carried out using the corrected factors rather than the non-corrected ones.

### C. Clinical Groups

AgeML offers the capability to examine the participant’s age deltas labeled into various clinical populations. It assesses the distributions of age deltas for each provided clinical population, displaying the mean and standard deviation for each group. Additionally, AgeML determines whether the delta distributions exhibit significant differences between groups using a paired t-Test, indicating whether p-values are significant to Bonferroni or FDR corrections for multiple comparisons. Effect sizes, confidence intervals, and the delta mean differences are also reported by AgeML. Similar to the factor correlation analysis, covariate correction can be applied to the deltas previous to calculating distributions for each clinical group.

AgeML also integrates a quality control step to check whether the age distributions of participants within each clinical group exhibit significant differences as training on a specific distribution and subsequently making predictions in a different one with a different age can potentially introduce biases [21]. This is particularly evident when considering young and old populations, as many phenotypes follow quadratic trends, as observed in brain volumes [32], [33] and cognitive performance [37]. Hence, training on a young population and applying the model to an older one can be problematic and susceptible to bias.

### D. Clinical Classification

In scenarios involving multiple organ-age systems, we represent **Δ** *≡ δ*_*i*_ as the collection of deltas for each subject, and use them as the input for a classifier designed to categorize participants into various clinical groups. Employing a cross-validation strategy, a logistic regressor is trained to differentiate between two specific labels. The sigmoid function is then given by 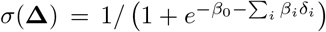, that after fitting the logistic regressor, the coefficient *β*_*i*_ accounts for the contribution of *δ*_*i*_ in the classification. The vector *β ≡ β*_*i*_, previously normalized between 0 and 1 by dividing each component by the maximum *β*_*i*_, is stored and printed to assess which organ delta participates with a higher relevance in the classification task.

## V. Results

### A. Reproducibility and Validation

To demonstrate that AgeML can replicate previous research findings, we chose to replicate the study by Tian et al. [1]. In this previous study a body age model is built along with 7 separate organ system age models for the cardiovascular, pulmonary, musculoskeletal, immune, renal, hepatic and metabolic systems. The different age models are trained on healthy participants with no known chronic disease. The feature set consists of the same 78 body phenotypes described in Data, and participants with any missing phenotypes are excluded from the analysis. Training is carried out using a Support Vector Regressor (SVR) with a 20-fold cross-validation scheme (with a linear kernel, a regularization parameter of 1.0 and a gap tolerance of 0.001). Comparisons of the results between AgeML (Ours) and the one in Tian et al. 2023 [1] are given in Table II.

**TABLE II.** Mean Absolute Error (MAE) and Pearson’s correlation (***ρ***) of SVR models for predicting the chronological ages of healthy controls using different physiological systems, comparing the work of Tian et al. 2023 [1] with ours.

| METRIC | GENDER | MODEL | BODY | CARDIOVASCULAR | PULMONARY | MUSCULOSKELETAL |
| --- | --- | --- | --- | --- | --- | --- |
| MAE | FEMALE | OURS | $3.86 \pm 0.21$ | $5.38 \pm 0.34$ | $5.34 \pm 0.27$ | $4.98 \pm 0.17$ |
| | | TIAN ET AL. 2023 [1] | $3.708 \pm 0.0015$ | $5.620 \pm 0.0016$ | $5.675 \pm 0.0008$ | $5.267 \pm 0.0010$ |
| | MALE | OURS | $4.61 \pm 0.21$ | $5.80 \pm 0.25$ | $5.55 \pm 0.23$ | $5.61 \pm 0.22$ |
| | | TIAN ET AL. 2023 [1] | $4.463 \pm 0.0016$ | $6.323 \pm 0.0011$ | $6.130 \pm 0.0008$ | $6.052 \pm 0.0012$ |
| $\rho$ | FEMALE | OURS | $0.753 \pm 0.034$ | $0.424 \pm 0.065$ | $0.443 \pm 0.064$ | $0.519 \pm 0.045$ |
| | | TIAN ET AL. 2023 [1] | $0.787 \pm 0.0002$ | $0.436 \pm 0.0004$ | $0.421 \pm 0.0003$ | $0.513 \pm 0.0002$ |
| | MALE | OURS | $0.638 \pm 0.033$ | $0.296 \pm 0.072$ | $0.382 \pm 0.051$ | $0.372 \pm 0.061$ |
| | | TIAN ET AL. 2023 [1] | $0.715 \pm 0.0002$ | $0.317 \pm 0.0004$ | $0.375 \pm 0.0003$ | $0.403 \pm 0.0003$ |

| METRIC | GENDER | MODEL | IMMUNE | RENAL | HEPATIC | METABOLIC |
| --- | --- | --- | --- | --- | --- | --- |
| MAE | FEMALE | OURS | $5.70 \pm 0.22$ | $5.16 \pm 0.24$ | $5.13 \pm 0.29$ | $4.97 \pm 0.24$ |
| | | TIAN ET AL. 2023 [1] | $5.965 \pm 0.0019$ | $5.220 \pm 0.0012$ | $5.466 \pm 0.0010$ | $5.302 \pm 0.0022$ |
| | MALE | OURS | $5.78 \pm 0.22$ | $5.41 \pm 0.31$ | $5.75 \pm 0.25$ | $5.85 \pm 0.24$ |
| | | TIAN ET AL. 2023 [1] | $6.330 \pm 0.0022$ | $5.830 \pm 0.0011$ | $6.294 \pm 0.0012$ | $6.514 \pm 0.0019$ |
| $\rho$ | FEMALE | OURS | $0.284 \pm 0.087$ | $0.475 \pm 0.069$ | $0.476 \pm 0.084$ | $0.524 \pm 0.064$ |
| | | TIAN ET AL. 2023 [1] | $0.303 \pm 0.0005$ | $0.532 \pm 0.0002$ | $0.451 \pm 0.0004$ | $0.498 \pm 0.0005$ |
| | MALE | OURS | $0.297 \pm 0.067$ | $0.437 \pm 0.058$ | $0.308 \pm 0.065$ | $0.247 \pm 0.065$ |
| | | TIAN ET AL. 2023 [1] | $0.301 \pm 0.0007$ | $0.453 \pm 0.0003$ | $0.322 \pm 0.0004$ | $0.230 \pm 0.0009$ |

\* the reported ± are standard deviations of the MAE and Pearson's correlation across CV splits for our work and for Tian et. al. 2023 [1] they are the standard deviations of the MAE and Pearson's correlation over all subjects repeating the modeling 10 times with different seeds for the CV scheme.

Tian et al. 2023 [1] cite the standard deviation of the Mean Absolute Error (MAE) and Pearson’s correlation (*ρ*) over all subjects repeating the modeling 10 times with different seeds for the CV scheme. AgeML calculates the standard deviation of the MAE and *ρ* across the CV splits. This measure is larger but more representative of variability when the model looks at new data compared to the standard deviation of repeating the same experiment 10 times with the same data but different CV splits. It is also computationally less expensive. AgeML results were within 2 of our standard deviations measures of the results reported in Tian et al. 2023 [1]. There are some minor discrepancies possibly attributed to the differences in sample sizes (Ours N=6,388; Tian’s N=28,589). Differences in sample size are due to differences in inclusion criteria, primarily our more stringent definition of chronic conditions.

We evaluated the performance of three different machine learning models (Ridge, Random Forest and Support Vector Machine) implemented in AgeML for age prediction across different organ systems. MAE and *ρ* for each organ system and model can be found in Supplementary File Table S.I. Across all organ systems, the error metrics for these models were generally within one standard deviation of each other, indicating similar performance. However, for the body model, which involved the most features, Ridge Regression demonstrated the best performance, while Random Forests exhibited the worst. We have also used AgeML with data from ADNI [38] to reproduce our previous work Garcia Condado et al. 2023 [7]. We obtain similar error metrics in the brain age modeling task when using structural and neuropsychological features as seen in the Supplementary File Table S.II. We also obtain similar distributions of the deltas for the three different clinical groups: Controls (CN), Mild Cognitive Impairment (MCI) and Alzheimer’s disease (AD) as seen in the Supplementary File Table S.III.

### B. Genetic Analysis

PRSs measure genetic risks for various diseases, and some diseases are closely related to specific organs, possibly influencing organ age. For example, it is expected that genetic predisposition to factors affecting the heart, such as cardiovascular diseases, coronary artery diseases, ischemic stroke, or hypertension, would lead to higher cardiovascular ages. Similarly, protective genetic factors that promote stronger bones with higher mineral density can lead to a reduction in musculoskeletal age. Here, and for the group of healthy participants, the correlation between age deltas for each organ and both Standard PRS and Enhanced PRS were compared, cf. Fig 2 shows statistically significant correlations between PRS and organ deltas (detailed correlation values and corresponding p-values are given in Supplementary File Table S.IV for Standard PRS and Table S.V for Enhanced PRS). Very remarkably, for the results corresponding to Enhanced PRS, we found that many metabolic genetic propensities negatively impact metabolic age, and in contrast, a protective effect of glomerular filtration. The differences in statistical significance between Standard PRS and Enhanced PRS may be due to the lower number of participants available for analysis in the latter.

**Fig. 2.**
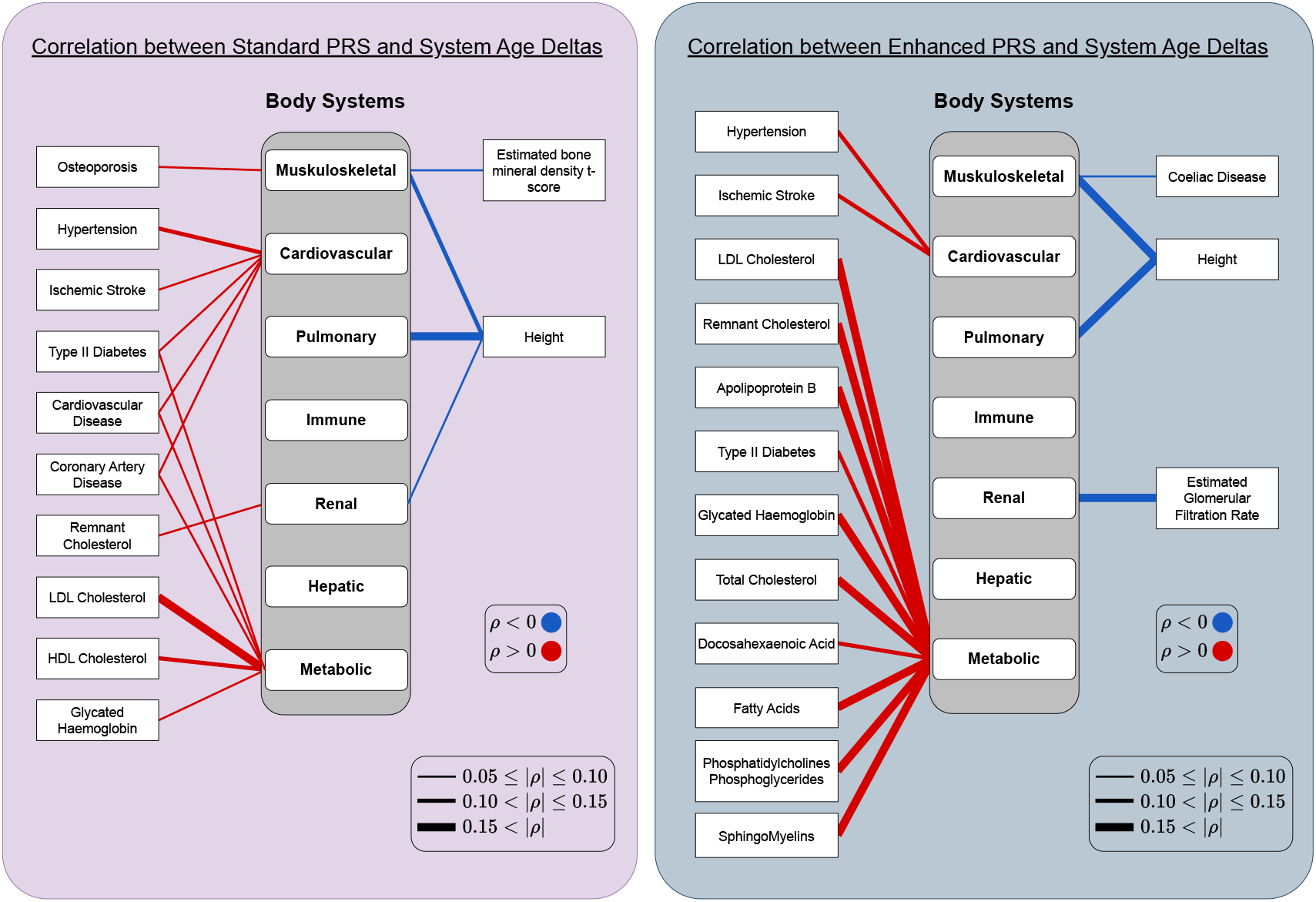
Factor correlation analysis between Polygenetic Risk Scores (PRS) and system-specific age deltas. Each link represents a statistical significant Bonferroni-corrected Pearson’s correlation (***p <* 0.05**) and only those with |***ρ***| ***>* 0.05** are plotted. The width of the link indicates the strength of the correlation. Red links denote positive correlations, which are associated with higher age deltas or accelerated aging. Blue links, on the other hand, represent negative correlations, suggesting protective or age-slowing effects. The left purple panel shows results for Standard PRSs (6,323 participants), while the right gray panel does it for Enhanced PRSs (1,234 participants). Note that several PRS labels are shared across panels, reassuring the significance of the correlations. All the Pearson’s correlation values and their associated p-values can be found in the code repository.

### C. Clinical data

Deltas for each organ systems were calculated for all participants. The average deltas for each clinical group and organ were calculated, and then compared across groups (Table III). The significance of these differences indicates that the participants belonging to clinical groups related to specific organs exhibit significant variations as compared to healthy controls. For example, individuals in the clinical category of circulatory diseases show a higher delta for the cardiovascular system compared to healthy controls. Detailed information about the differences in average deltas and their corresponding p-values, effect sizes, and confidence intervals between clinical groups and organ systems can be found in the Supplementary File Tables S.VI and S.VII, respectively.

**TABLE III.** Disease categories that have a statistically significant difference in their age delta compared to healthy controls, for different systems (Bonferroni corrected, ***p <* 0.05**).

| SYSTEM DELTA | CLINICAL CATEGORY DIFFERENT TO CN |  |  |  |  |
| --- | --- | --- | --- | --- | --- |
| CARDIOVASCULAR | CIR | MBD | MUS | RSP | NEO |
| PULMONARY | CIR | MBD | RSP | NEO |  |
| METABOLIC | MUS | NEO |  |  |  |
| IMMUNE | MBD |  |  |  |  |

CN: Healthy Controls, CIR: Circulatory system disease, MBD: neurodevelopmental disorders, MUS: Musculoskeletal system disorders, RSP: respiratory system diseases, NEO: neoplasms, BLD: blood diseases

Logistic regressor classifiers were trained to differentiate between healthy controls and each clinical category based on all organ deltas as input. Fig 3 displays the logistic regressor weights, which have been normalized for each case. Similar to the differences observed in average deltas for organ systems in specific clinical categories, the most discriminant features between healthy controls and clinical categories related to specific organs are the deltas for those respective organs. Thus for instance, when classifying between healthy controls and cardiovascular disease, the most discriminant feature is the cardiovascular delta. To avoid biases in predictions, the majority class was under-sampled to match the size of the minority class.

**Fig. 3.**
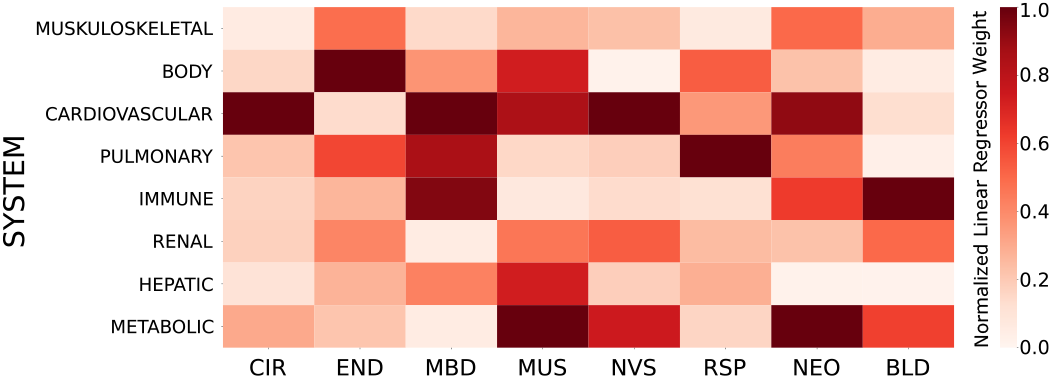
Normalized Logistic Regressor weights of system-specific deltas for classification between healthy controls versus a specific disease category. The highest values for each column indicate the most relevant system-specific delta used by the Logistic Regressor for classifying healthy controls against the column disease category. Normalization is done per column (classifier), dividing the absolute values by the absolute highest one. Thus, the cells with the value 1.0 correspond to the absolute highest weight per column. In the situation of class imbalance, the majority class was undersampled to match the minority class occurrences.

## VI. Discussion

The first step towards standardization is showing the reproducibility of previous findings. In this context, we have replicated the study presented by Tian et al. [1]. AgeML facilitates standardization by providing a tool designed for use with explicitly defined pipelines. Moreover, the reporting of standardized metrics by AgeML constitutes a crucial issue for usability. AgeML aims to establish the minimum guidelines for clear communication of training outcomes.

Previous research like BASE [22] focused on brain age modeling has shown the need for standardization and reproducibility. While BASE is tailored to brain age modeling, it highlights the significance of integrating standards into the field. AgeML not only replicates many of the benchmarks established in BASE with respect to metric reporting but also extends it further by enhancing ease of access, embracing open-source development, and demonstrating applicability across various types of clinical or organ data.

When exploring the performance of various machine learning models for age prediction, extending our analysis beyond Support Vector Regression (SVR) to include Ridge Regression and Random Forests, the results demonstrated that, across most organ systems, the error metrics for these models were within one standard deviation of each other. This suggests that the choice of model had minimal impact on predictive accuracy in these cases. In the body model, which featured the highest number of features, Ridge Regression outperformed the other models, while Random Forests performed the worst. This highlights the importance of considering the complexity and feature density of datasets when selecting a machine learning model. Although we applied hyperparameter tuning, performance improvements were minimal, consistent with findings from previous research [1]. These results suggest that while hyperparameter tuning and model selection remain important considerations, the inherent characteristics of the dataset may play a more significant role in determining model success. Future work should continue exploring model performance across a wider variety of datasets to further refine these predictions.

A potential limitation of AgeML is the risk of overfitting when constructing biological age models using only healthy cohorts and applying these models to both healthy and patient populations. This could affect the model’s generalizability and the validity of comparisons between groups. It is an open discussion in the field whether models should be trained on a healthy population or a more diverse dataset representative of the general population. To address this, we propose two strategies. First, models can be trained on a broader population that includes both healthy and patient groups and is representative of the general population under study. Researchers must ensure this training set is separate from the test set. Second, AgeML incorporates regularization techniques and cross-validation to mitigate overfitting during model construction. Both approaches can be applied simultaneously for more robust model performance. While these measures help reduce overfitting, users should interpret results carefully when dealing with heterogeneous datasets. We acknowledge this limitation and suggest future improvements, including the integration of pretrained models derived from larger, more diverse populations, which could enhance generalizability across various cohorts.

We have shown that genetic information, specifically Polygenic Risk Scores (PRS), are associated with distinct forms of organ aging, aligning with prior GWAS studies [9]. This is of critical importance for quantifying the extent to which an individual’s organ age-variations can be attributed to genetics. Moreover, it paves the road for exploring therapeutic targets aimed at decelerating aging [9]. A logical continuation of this work is to develop organ-specific PRS based on genomic data to assess genetic predisposition to either significantly rejuvenated or minimally aged organs as a measure of aging resilience. Efforts in this direction are already in progress, notably in the context of brain age PRS [39].

The Standard PRS is derived from external datasets, ensuring broader applicability across populations but potentially sacrificing some degree of accuracy. In contrast, the Enhanced PRS incorporates additional training data from the UK Biobank itself, improving prediction accuracy but limiting its generalizability outside the UK Biobank population due to cohort-specific overfitting concerns. One practical implication of these differences lies in the sample size disparity: the Enhanced PRS (1,234 participants) may offer increased predictive power due to the enriched dataset, but the smaller sample size relative to the Standard PRS (6,323 participants) reduces its generalizability and increases the risk of bias. The smaller sample size of the Enhanced PRS could result in lower statistical significance, limiting the robustness of conclusions drawn from it. On the other hand, the Enhanced PRS may be less prone to biases from external datasets, offering a more tailored risk prediction in the UK Biobank cohort. To mitigate overfitting, we encourage the use of cross-validation and suggest that users carefully interpret results depending on the context in which these models are applied.

As expected the age deltas of each organ systems vary among clinical populations. This variation is particularly evident in the deltas associated with the cardiovascular system and its corresponding clinical category of cardiovascular disease. Similar to previous work [1], our results confirm that while large age deltas are not exclusive to any single disease, distinct patterns of organ deltas are associated with each specific disease.

The practical integration of age modeling in clinical environments is currently in a conceptual phase. An initial proposition involves using age deltas as a flag, where during routine analyses, these deltas are computed, and an alert is triggered if any of the calculated deltas are high (previously determined a normal or standardized distribution of deltas). Although organ deltas lack specificity to particular diseases (eg., patients with Parkinson’s, Alzheimer’s, or depression might exhibit an elevated brain delta), they can alert clinicians to potential deviations from healthy aging normative patterns. By computing deltas for various organs and achieving a multimodal perspective, it is also feasible to mitigate the issue of non-specificity, thereby enhancing the feasibility of age deltas in clinical settings.

AgeML is a first approach to setting a standard in the community and a minimal software tool for basic age modeling. The AgeML project will continue by integrating more features. Currently AgeML supports only tabular data; however, future plans aim to incorporate additional data modalities, including imaging data (e.g., MRI or CT scans) routinely used in medical settings, as well as time series data, such as electrocardiograms (ECG). The next step involves integrating foundational models that have been trained on extensive datasets, such as the UKBiobank. Following that, the strategy includes providing transfer learning capabilities to retrain models with local datasets. In medical contexts, the demand for interpretability is paramount and should be addressed by using for example Shap values [40]. The factor correlation pipeline could be upgraded to introduce causal analysis. Ultimately, it is anticipated that community-driven efforts will emerge, aiming to incorporate a broader range of desired features and improvements.

## VII. Conclusion

We have shown that AgeML is capable to replicate prior studies in the field, thereby paving the way for standardization of age modeling strategies. We have proven the AgeML efficacy by investigating the interplay between age-related organ variations and genetics, and in particular, by assessing the correlation between Polygenic Risk Scores and specific organ deltas. Consistent with previous work, statistical significant differences in organ deltas occur between healthy controls and clinical populations, especially when focusing on organ-systems related to specific diseases.

The ultimate objective of our study extends beyond merely developing a tool; it aims to trigger a community of developers who contribute new features and help lead towards age-modeling standardization. The major goal of AgeML is to standardize procedures, reduce the barriers to entry for age modeling, and guarantee the reproducibility of research findings. The AgeML project is Open-Source, designed to foster a welcoming environment and a community to work together to improve and validate existing methodologies. We are actively inviting new developers interested in contributing to the growth and expansion of the package.

## Acknolwedgments

This research has been conducted using the UKBiobank Resource under Application Number 79105.

## Notes

The project that gave rise to these results received the support of a fellowship from “la Caixa” Foundation (ID 100010434) to JGC. The fellowship code is LCF/BQ/DI21/11860030. JMC is funded by Ikerbasque: The Basque Foundation for Science, the Department of Economic Development and Infrastructure of the Basque Country (Elkartek Program, grant KK-2021-00009), and the Health Department of the Basque Country (grants 2022111031 and 2023111002). AE is supported by Ikerbasque, the Basque Foundation for Science, and the Spanish Ministry of Science and Innovation, grant RYC2021-032390-I.

https://github.com/compneurobilbao/ageml

### Competing Interest Statement

The authors have declared no competing interest.

### Summary of Updates

Revision of the manuscript. Abstract updated, better wording.

